# Identifying Context-specific Network Features for CRISPR-Cas9 Targeting Efficiency Using Accurate and Interpretable Deep Neural Network

**DOI:** 10.1101/505602

**Authors:** Qiao Liu, Di He, Lei Xie

**Affiliations:** Department of Computer Science, Hunter College, The City University of New York; Ph.D. Program in Computer Science, The Graduate Center, The City University of New York; Ph.D. Program in Biochemistry, The Graduate Center, The City University of New York

**Author notes:** Corresponding author Lei Xie, Ph.D. Program in Biology, Biochemistry, & Computer Science, and Department of Computer Science, Hunter College, The City University of New York 695 Park Avenue, New York City, NY 10065, U.S.A.

## Abstract

CRISPR-Cas9 is a powerful genome editing tool, whose efficiency and safety depends on the selection of single-guide RNA (sgRNA). Machine learning has been applied to optimize sgRNA selection, but several challenges remain. The performance of predictive model is limited by the amount of available data in many cell lines, ignorance of gene network function and its variable effect on phenotype, and elusive biological interpretation of computational models. We develop an accurate and interpretable deep learning model SeqCrispr to address these problems. In benchmark studies, SeqCrispr outperforms state-of-the-art algorithms and improves the prediction accuracy when applied to small sample size cell lines. Furthermore, we find that gene context-specific network properties are critical for the prediction accuracy in addition to the last three nucleotides in sgRNA 3’end. Our findings will bolster developing more accurate predictive models of CRISPR-Cas9 across wide spectrum of biological conditions as well as efficient and safe gene therapy.

## Introduction

The clustered regularly interspaced short palindromic repeats (CRISPR)-Cas9 genome engineering is a powerful tool for modifying specific genome DNA targets ^1, 2^. It was easily adopted to different biological systems, like plants, animals and even human cells, thus widely used in many applications. However, several optimizations are still necessary to improve this technique’s efficiency and specificity.

The targeting process in CRISPR-cas9 system has three fundamental requirements for its success, specifically for *S. pyogenes* Cas9 ^3, 4^. First, 20 nucleotide single-guide RNA (sgRNA) needs to be complementary with targeting genome sequence. Second, a Protospacer Adjacent Motif (PAM) needs to be located upstream of the target site ^5^. Finally, off-target effect, which is caused by other genome sequences similar to the targeting sequence, needs to be minimized ^6^. These are necessary for a powerful system, but are not sufficient. Tens or hundreds of sgRNAs can be chosen from to knock out or modify a target gene, but not all of them are ideal. Different factors were proposed to affect sgRNA on-target efficiency and specificity. For instance, the first step of sgRNA targeting is the unwinding of targeted genome dsDNA ^7^. The strength of target-site DNA double strands binding is a determinant for the unwinding rate and thus has impact on sgRNA targeting efficiency. DNase sensitivity can indicate the chromatin coverage and accessibility of target sites ^8, 9^. Open chromatin sites may promote sgRNA binding efficiency due to their high accessibility. Epigenetic features, such as transcription factor binding and methylation on target DNA strand, could also affect sgRNA targeting efficiency ^10, 11^. It was believed that successful design of sgRNAs could save hours and days work as well as the cost on experimental reagents ^12^. Thus, determination of critical biological features for the on-target efficiency and computational prediction of sgRNA on-target efficiency will significantly facilitate the success of CRISPR-Cas9 experiments.

Predicting sgRNA targeting efficiency is still a challenging problem. Especially, sgRNA targeting efficiencies are different when experiments are not conducted in the same cell line. Differences between cell lines can result from re-wired biological network, varied basal gene expressions, different epigenetic features, and genome structural variation such as copy number differences. Many system-level omics data are now available benefiting from wide utilization of next generation sequencing ^13, 14^. Besides, genome-wide screening technique that combined the CRISPR-Cas9 technique and next generation sequencing methods made large-scale CRISPR-Cas9 data available ^15^. With these large-scale datasets, it becomes feasible to integrate gene copy number variation, gene expression, epigenome information, and sequencing data to develop machine learning models. Up to date, several deep learning models have achieved considerable success ^16, 17^.

In spite of these progresses, several critical issues remain in the understanding and the prediction of CRISPR-Cas9 efficiency. First, even though large-scale dataset is now available, not all of cell lines have enough data for training accurate machine learning models. Second, the state-of-the-art deep learning model is a black-box. Their biological interpretation is still elusive. It is not clear what the most important biological features are to determine the on-target efficiency. Last, because of the lack of rational methods comparing sgRNAs efficiencies targeting on different genes with distinct functions and essentialities, compromises had to be made. Few existing methods take the context-specific network functional difference of targeting genes into account. Since cellular response to gene editing is a systematic behavior, global network properties of a gene may play an important role. However, they have not been explored in the machine learning.

In this paper, we have made several contributions to address these problems. We have developed a new deep learning model SeqCrispr. SeqCripr have five unique features. First, it uses a novel representation of nucleotides. Second, it for the first time incorporates context-specific network features of gene into the model. Third, it combines two most successful deep learning architectures: recurrent neuron network (RNN) and convolutional neuron network (CNN). Fourth, it uses transfer learning to train predictive models for the cell lines that have few samples. Finally, it implements a universal feature ranking algorithm for the deep learning to determine the importance of biological features responsible for the CRISPR-Cas9 on-target efficiency. In the benchmark study, SeqCrispr outperforms the state-of-the-art algorithms ^16, 17^ as well as significantly improves the prediction accuracy when applied to the cell line with a small sample size. Besides, we find that the context-specific network feature is critical for the CRISPR-Cas9 on-target efficiency, in addition that the identity of last three nucleotides in sgRNA 3’end is the most important feature of sgRNA. Our results may shed new light into developing more accurate and robust machine learning models for selecting efficient sgRNA across wide spectrum of biological conditions as well as has implications on developing efficient and safe gene therapy using CRISPR-Cas9 technique.

## Results

### Overview of SeqCrispr model for CRISPR-Cas9 efficiency prediction

The proposed SeqCrispr model for the prediction of CRISPR-Cas9 efficiency has four major components, as shown in Figure 1. The first component is a nucleotide embedding layer that is inspired by the word2vec technique ^18^. As described in Methods part, unsupervised representation learning for vector representation of 3mers was conducted with whole genome exons sequences. The vector representations of 3mers in the embedding layer of SeqCrispr were initialized with that learned with unsupervised representation learning and were also fine-tuned later. In addition to the 3mer representation, local biological features such as DNA-sgRNA binding melting temperature, DNase peak, CTCF peak, RRBS peak, and H3K4me3 peak as well as global gene network property that is derived from the gene neighbor connection and their expression values are also used as input features. The second component is a hybridization of recurrent neuron network (RNN) and convolutional neuron network (CNN) for the feature engineering of the sgRNA. The prior one is well known for its good performance on analyzing sequential data, like natural language processing ^19^. The performance of later one is superior on image data processing, but evidences also show CNN’s potential on sequential data processing, such as sentence classification ^20^. Thus these two models could be complementary for analyzing sgRNA efficiency considering that the primary feature of CRISPR-Cas9, sgRNA sequence, is also sequential. The third component is a fully connected deep neural network that combine the sequence features of the sgRNA with the local and global biological features of targeting gene as the input. The fourth compound is added to rank the importance of sequence and biological features using input perturbation method.

**Figure 1.**
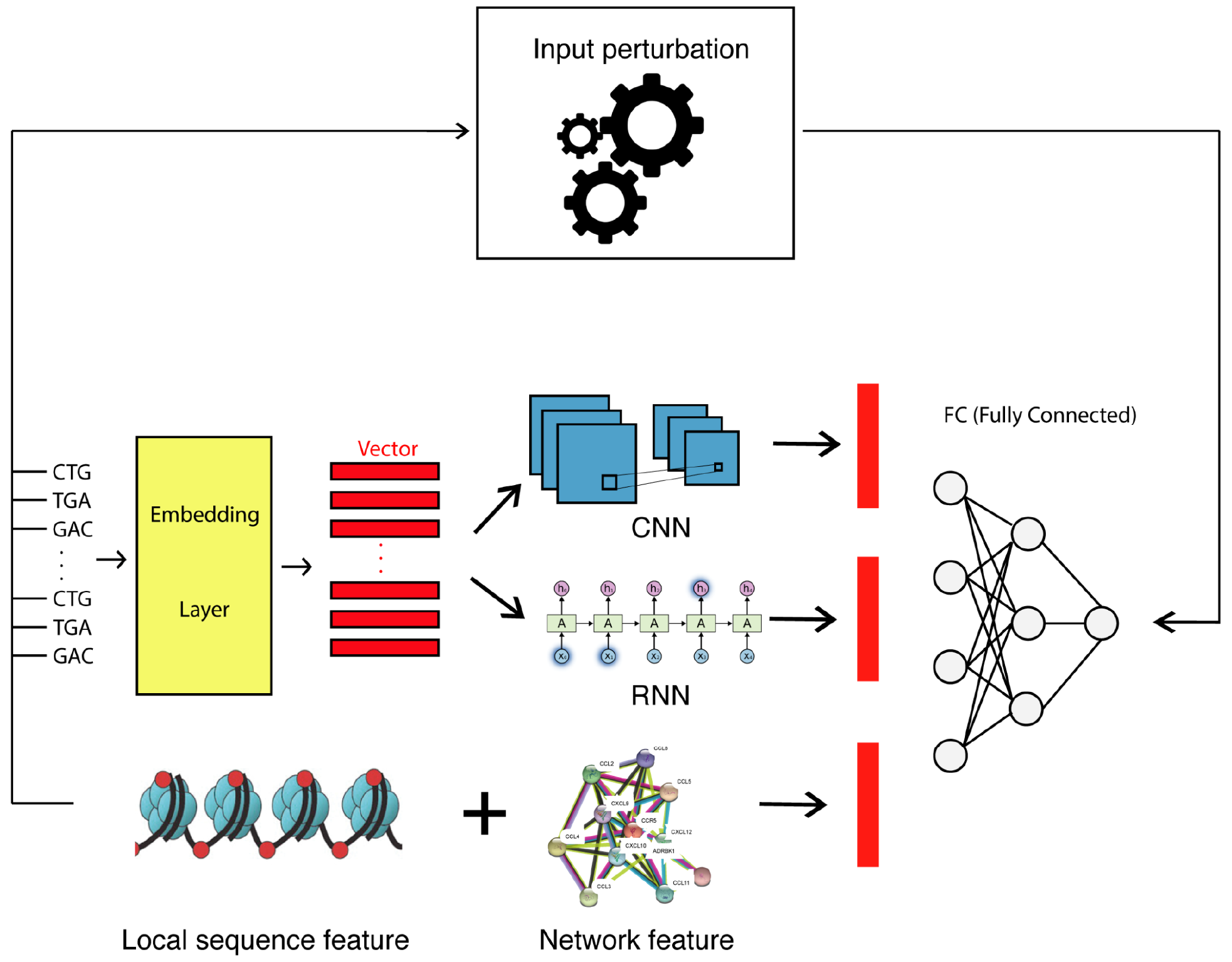
Schematic representation of SeqCrispr Model. This model includes four components: 1) Embedding layer, 2) Convolutional neural network and recurrent neural network layer, 3) Fully connected layer, and 4) Input perturbation layer.

### SeqCrispr hybrid model is more accurate than state-of-the-art deep learning models

We compared the performance of SeqCrispr hybrid model with random forest, boosted regression trees, SeqCrispr RNN only, and SeqCrispr CNN only. The SeqCrispr hybrid model clearly outperformed other models (Figure 2A). The evaluation metric is spearman correlation of predicted on-target efficiency and ground truth on-target efficiency. The performance of SeqCrispr hybrid model is higher than conventional machine learning models by 3%-18%. It can perform better than SeqCrispr CNN by 4.7%-11.6% and SeqCrispr RNN by 0%-6.7%. Thus we used SeqCrispr hybrid model to perform follow-on tests and comparisons.

**Figure 2.**
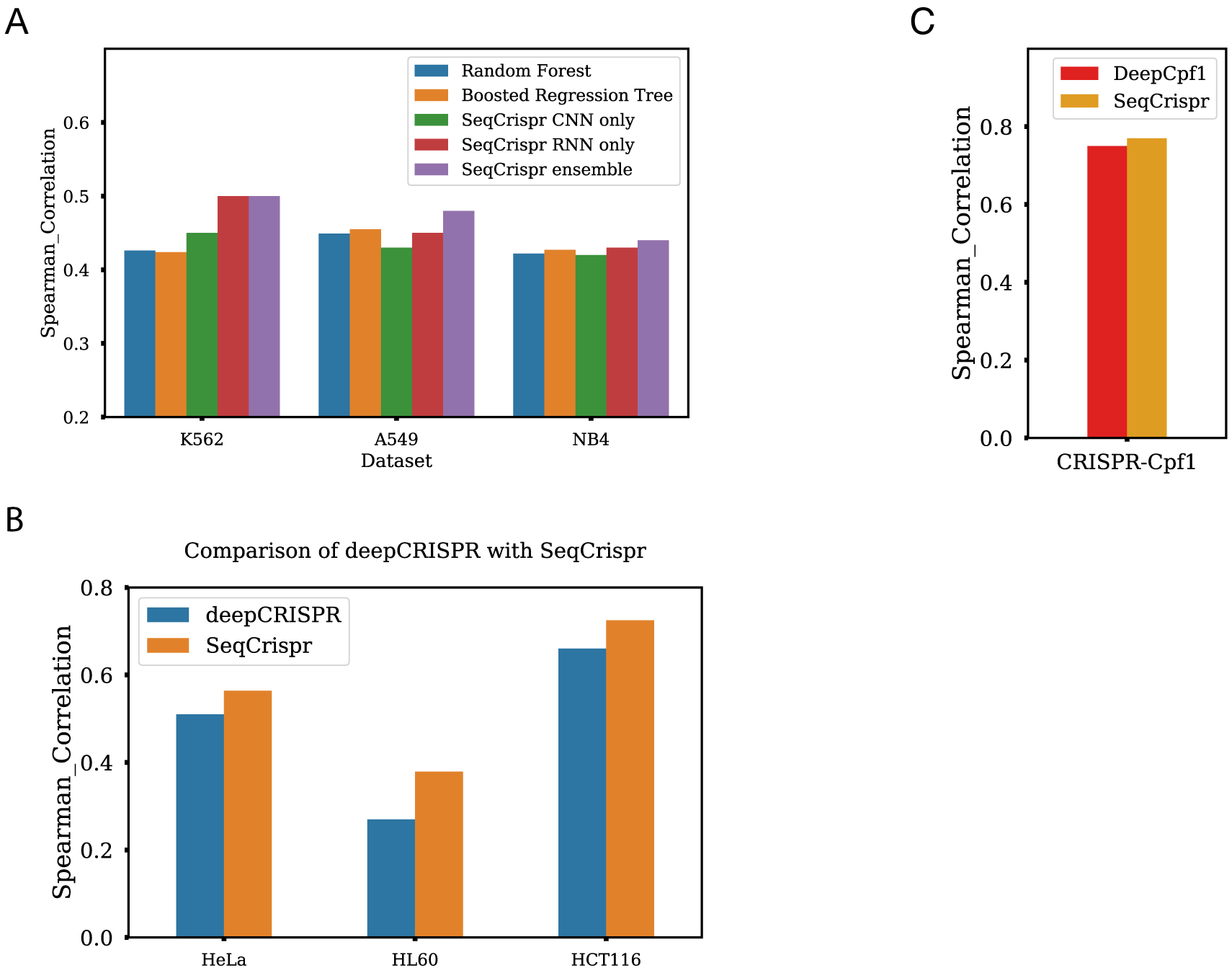
SeqCrispr hybrid model can get comparable performance with or higher performance than other state-of-art deep learning models. A) Comparison of random forest model, boosted regression tree model, SeqCrispr models performance on dataset collected in K562, A549 and NB4 cell lines. Deep learning outperforms conventional machine learning model. B) Comparison of SeqCrispr model and deepCRISPR performance on dataset collected in each of HeLa, HL60 and HCT116 cell lines. C) Comparison of SeqCrispr model and DeepCpf1 model performances with dataset collected with CRISPR-Cpf1 experiment.

We further compared the SeqCrispr with the state-of-the-art deep learning models that were trained with other datasets. We evaluated the SeqCrispr model in two different scenarios using spearman correlation metric. 1) Pre-selected negative selection dataset. It was used to train deepCRISPR model ^16^. The original dataset has ~15K sgRNAs data from four cell lines: HeLa, HL60, HCT116 and HEK293T. However, gene expression profile data is not available in HEK293T cell line, so the gene global functional property couldn’t be computed. We only studied the remaining three cell lines data. We combined randomly selected 80% data from each cell line to train the SeqCrispr model and then tested the left 20% data in each cell line. The spearman correlation in each of the three cell line is all better than the published performance of deepCRISPR model (Figure 2B). 2) CRISPR-Cpf1dataset ^17^. Deep learning model had better performance on studying CRISPR-Cpf1 data. SeqCrispr model spearman correlation performance on this dataset is around 0.77. As a comparison, published model DeepCpf1 used in Kim et. al could get similar spearman correlation, ~0.75 (Figure 2C). Since only the sgRNA sequence information is available, the model input only includes sgRNA primary sequence.

In summary, our implemented deep learning models SeqCrispr has comparable performance with or higher performance than the current state-of-art deep learning models. It is noted that the word2vec embedding of 3-mer nucleotides outperforms the corresponding one-hot encoding representation that has been widely used by the state-of-the-art methods (Supplementary material Table S1). Interestingly, word2vec embedding with Hilbert-curve filling may have advantage over vertical stacking (Table S1) ^21^. We also noted that the performances of machine learning models were also determined by the quality of experimental data. Besides, the SeqCrispr model performance on CRISPR-Cpf1 experiment is higher than that on CRISPR-Cas9 experiment. This might be attributed to lower off-target effect in CRISPR-Cpf1 system ^22, 23^.

### Learned information are transferable between cell lines

We further investigated the generality and transferability of the SeqCrispr model. We firstly learned model with one cell line’s data and then tested the performances of the model on another cell line’s data. The spearman correlation ranges in 0.24-0.37 without any posterior weights fine-tuning (Table I). This indicates that the information learned with data of one cell line is transferable to other cell lines. Next, we tested how transferable each part of SeqCrispr model is. The first model is trained with the combined dataset from K562, A549 and NB4 cell lines. Transfer learning was conducted on the combined dataset from HeLa, HL60 and HCT116 cell lines. We froze the weights in the first few layers and fine-tuned the remaining layers’ weights, respectively. We showed that the performance would drop more when more layers were frozen (Figure 3A). For the following analysis, we chose to only freeze the embedding layer, a trade-off between performance and number of trainable parameters. Then we performed transfer learning with new data in different sample sizes (Figure 4B). We found fine tuning the unfrozen weights of SeqCrispr model with more unstudied data could still boost model performance. Thus, when training data is limited, transfer learning method can improve predictive model performance (Figure 4B). It is also worth noting that the performance of the trained model by the regular training method could catch up with that of the model trained by the transfer learning method when sample size is large enough. Thus when a cell line data sample size is small, a model learned with another cell line data has transferable information and initializing parameters with that of trained model could improve new model’s prediction performance in this cell line. However, more data is still necessary to deliver cell line specific information to the model to boost performance.

**Table I.** performances of the model training the dataset of cell line in each row and testing on the dataset of cell line in each column.

|  | K562 | A549 | NB4 |
| --- | --- | --- | --- |
| K562 | 0.50 | 0.325 | 0.252 |
| A549 | 0.282 | 0.48 | 0.36 |
| NB4 | 0.24 | 0.37 | 0.44 |

**Figure 3).**
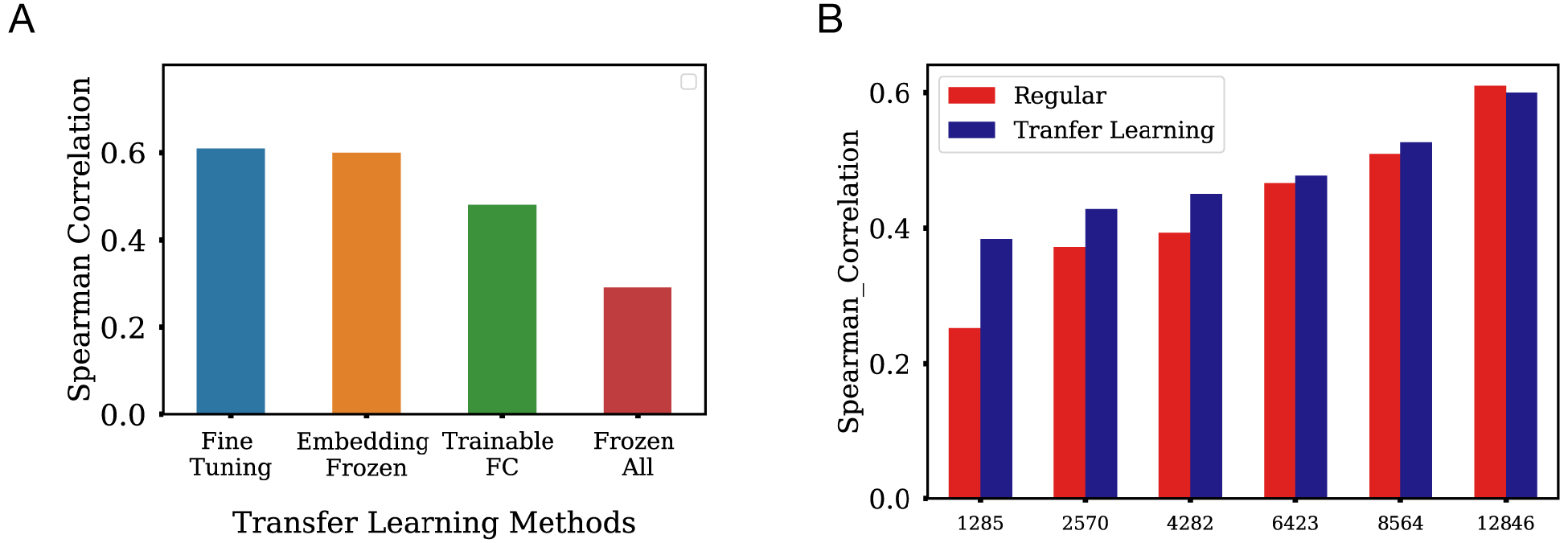
Transfer learning performances of SeqCrispr model that is trained with combined dataset from K562, A549 and NB4 cell lines and then fine-tuned on combined dataset from HL60, HCT116 and HeLa cell lines. A) Different layers’ weights are frozen and made untrainable. Fine tuning: All weights in the model are trainable. Embedding frozen: Only the weights matrix in embedding layer are frozen and the remaining weights are trainable. Trainable FC: only the weights in the last fully connected layers are trainable and all the remaining weights are frozen. Frozen all: All the weights are frozen. B) Transfer learning was performed in Embedding frozen way. Transfer learning: different sizes of training data from HL60, HCT116 and HeLa cell lines are used to fine-tune the transferred model and same test data is used. Regular: different sizes of training data from HL60, HCT116 and HeLa cell lines are used to train a new model with randomly initialized weights and tested with the same test dataset.

### Context-specific gene network property boosts models performance and is one of the most important features responsible for CRISPR-Cas9 on-target efficiency

In CRISPR-Cas9 system, sgRNA is the key to find target genome location by matching sgRNA sequence with target genome complementary sequence. The sgRNA on-target efficiency can not only affected by sgRNA primary sequence, but also by other related biological features. The data we used in this study are from CRISPR-Cas9 negative selection experiment in three cell lines (K562, A549 and NB4). These cell lines were selected because their biological features, including epigenetic features, gene expression, and copy number, are available. In a negative selection experiment, a sgRNA pool is firstly designed. Cells are then transduced by sgRNAs and sgRNAs targeting on essential genes cause cell death or inhibit cell proliferation ^24^. Next generation sequencing technology is used to detect sgRNA counts in control group and experimental group. Log2fc is calculated afterwards. A more negative log2fc indicates a higher sgRNA targeting efficiency. Low sgRNA targeting efficiency is due to either low sgRNA targeting efficiency or low gene essentiality.

We chose several biological features as our models inputs. They include local sequence properties that affect the accessibility of target genome, and quantify sgRNA and DNA binding efficiency. These included amino acid cutting position, RNA-DNA binding melting temperature and several epigenetic features. 1) Amino acid cutting position. A mutation close to 5’ end of the gene is more likely to cause a structural change on expressed protein because transcript sequence locating downstream of the mutation position will be changed. 2) sgRNA-DNA melting temperature. Higher sgRNA-DNA binding melting temperature indicates higher binding efficiency. 3) We included multiple epigenetic features, including DNase, CTCF binding, H3K4me3 and RRBS. Genome regions that are in open chromatin are more sensitive to DNase cleavage ^25^. Target DNA sequence in open chromatin region could be more accessible by sgRNA. CTCF binding was shown to regulate high order chromatin structure ^26^. The methylation of H3K4 is also an important epigenetic mark associated with transcription factors binding and chromatin remodeling ^27^. Even though some studies argued that DNA methylation had no significant effect on Cas9 cutting on target genome which can perfectly match with sgRNA ^11^, target gene methylation positions were still found to be negatively correlated with DNase hypersensitive sites ^10^. Thus we also included this feature.

We have also included copy number variance. Data has shown that sgRNAs targeting on DNA sequence with higher copy number led to higher anti-proliferation effect because they could have multiple targeting sequences on genome ^28^. Moreover, for the first time, we included context-specific gene network property that was derived from tissue-specific gene networks and gene expression profile. The sgRNAs targeting on non-essential or less-essential genes tend to show more positive log2fc because knocking out these genes will not inhibit cell proliferation. Other works have empirically filtered out the sgRNAs targeting on non-essential or less-essential genes, which dramatically decrease the available sample size ^12, 16^. Previous study had shown that unsupervised representation learning with all available sgRNAs targeting whole genome could result in better sgRNA vector representations and ameliorate the model’s performance ^16^. We believe that it is necessary to use more available data including sgRNA data targeting on some non-essential or less-essential genes. A less-essential gene in one cell line could be essential in another cell line. We included a quantitative score to represent context-specific gene network property. We compared three possible ways, gene centrality in a systematic gene network, gene expression, and a score calculated with both information ^29^. The calculated context-specific gene network property score showed best negative Pearson correlation with average log2fc for each gene (Table II), followed by gene centrality. It indicates that the gene network property of a gene is critical for the on-target efficiency, which has not been taken into account before.

**Table II.** Pearson correlation of average log2fc with gene expression, gene centrality and context-specific gene network property score.

|  | K562 | A549 | NB4 |
| --- | --- | --- | --- |
| Gene expression | -0.119 | -0.12 | -0.13 |
| Gene centrality | -0.242 | -0.29 | -0.246 |
| Gene network property score | -0.337 | -0.365 | -0.334 |

To determine the features importance in deep learning models, we implemented an input perturbation method which can be used to study any deep learning model features importance. With negative selection experiment data (K562, A549 and NB4 cell lines) (Figure 4A), we showed that gene network property could boost model performance. Besides, we observed that 1) the feature importance of gene network property is either ranked the first or second. 2) pos_18, which is the 18^th^ -20^th^ nucleotides in sgRNA, contributes significantly in these models. 3) The copy number variation is the second most important biological features. 4) The epigenetics features have lower importance than sequence features (Figure 4B, 4C and 4D). It is also worth to note that PAM sequence’s feature importance is low because all studied sgRNAs had optimized PAM sequence.

**Figure 4.**
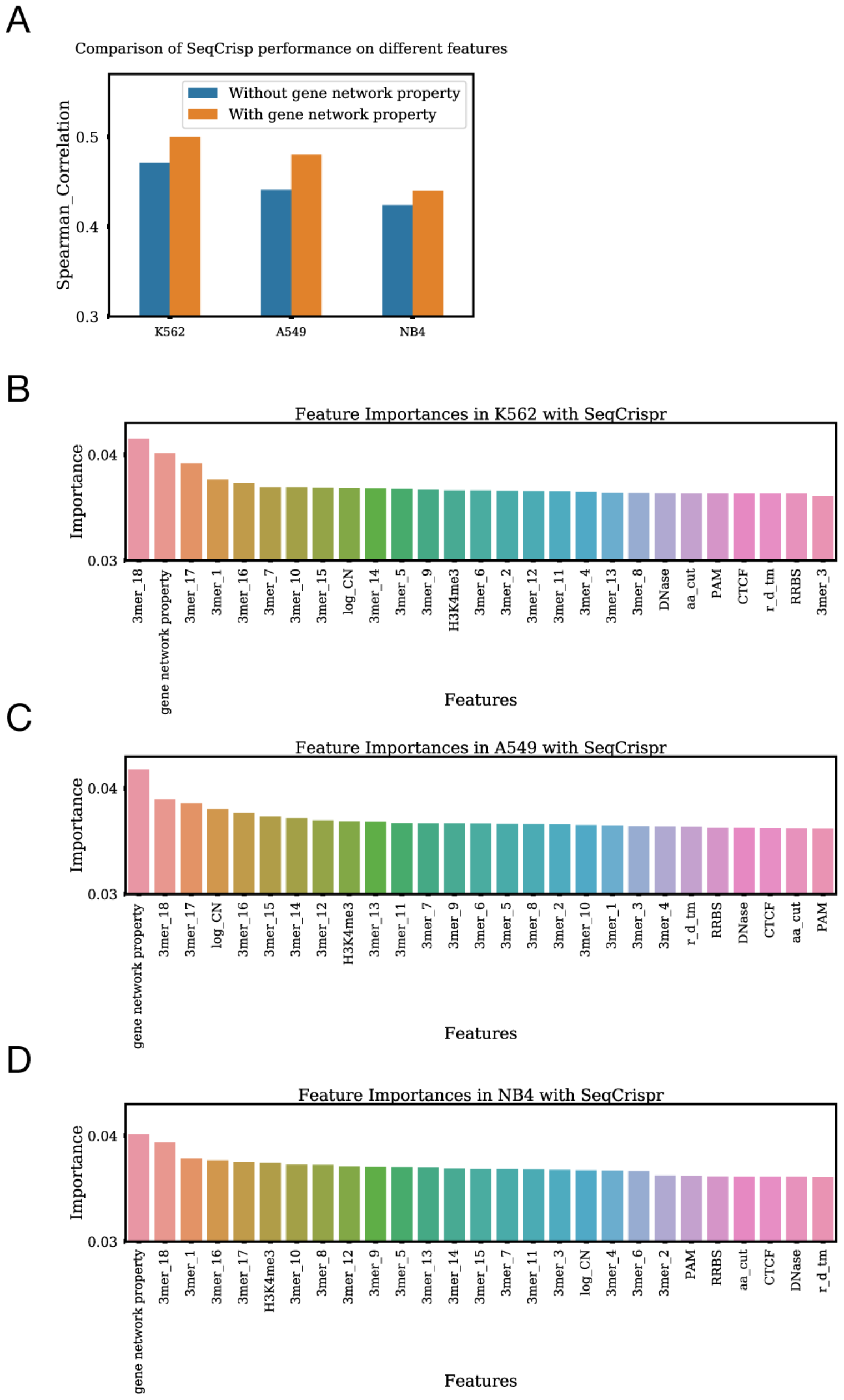
Features importance study for SeqCrispr hybrid model. A) SeqCrispr hybrid model was trained and tested on data with and without gene global functionality property score as a feature. The spearman correlation metrics were compared. B), C) and D) features importance of SeqCrispr models trained with dataset in B) K562 cell line C) A549 cell line and D) NB4 cell line are calculated with input perturbation method.

Similar results were also observed in the Random Forest model. We selected random forest models for three reasons. 1) As one of the state of art models, it is robust and resistant for overtraining ^30^. 2) It can be used to study features importance ^31^. 3) Both random forest and boosted regression trees model showed superior performance in previous studies. They have similar performance on the three datasets in this study. We drew the following conclusions (Figure 5). Including gene network property boosted models performance. The other important feature is pos_18 3mers, which is the 18^th^ -20^th^ nucleotides in sgRNA. This observation is supported by two experimental discoveries. Firstly, unwinding of target site dsDNA starts from the 3’ end of sgRNA ^7^. It showed that the unwinding process is critical for an efficient sgRNA targeting. Secondly, the cutting site of Cas9 endonuclease located in the 18^th^ nucleotide ^3, 32^. The identity of this nucleotide is a key factor for endonuclease cleavage on DNA genome. The consistent ranking of feature importance from different algorithms suggests that the determined biological features are model agnostic and could be biological meaningful.

**Figure 5.**
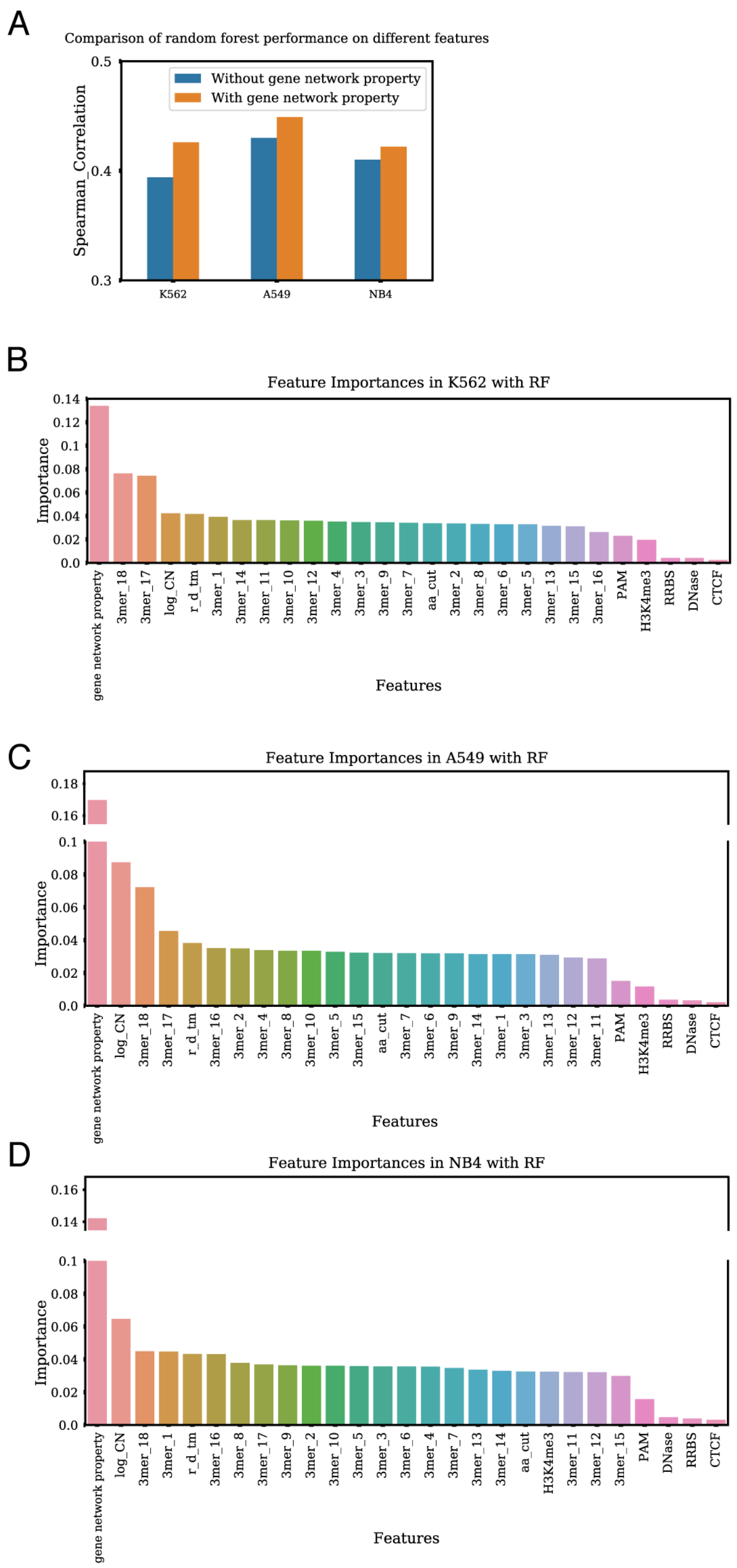
Features importance study for Random Forest model. A) Random Forest model was trained and tested on data with and without gene essentiality score as a feature. The spearman correlation metrics were compared. B), C) and D) feature importance attained with random forest model with data in B) K562 cell line C) A549 cell line and D) NB4 cell line.

## Discussion

Recently, large scale CRISPR-Cas9 experiments dataset were generated through the combination of next generation sequencing techniques. Machine learning models, particularly deep learning, could be used to study these datasets. In positive selection experiment, a selective reagent is applied to the sgRNAs library pool targeted cell populations. Knocking out one or a small fraction of genes could make cells gain resistance and survive. Doench et. al had identified an algorithm for on-target efficiency prediction, named Rule Set II, with ~4000 sgRNAs targeting on 17 genes ^6^. To make efficiencies of sgRNAs that are targeting on different genes comparable, they ranked all the sgRNAs’ log2fc targeting on one gene and used the rank to represent sgRNAs efficiencies. They assumed that sgRNAs efficiencies in one gene follows a uniform distribution. In negative selection experiment, the amount of data available is much larger because they don’t have any requirement for targeted genes. sgRNA libraries are transduced to a cell population. The cells with essential genes targeted by inserted sgRNA would lose the ability to proliferate due to genes loss of function and then die out. The sgRNAs log2fc could still indicate its on-target efficiencies. However, these introduce more variability due to different gene function and cellular response. Previous works pre-selected negative selection experiment data. They have set a threshold and pre-selected genes that was predicted to be essential genes and then limited analysis on sgRNAs targeting on these essential genes. All of existing methods ignore the variability and context-specific nature of gene essentiality. The impact of knocking-down/out a gene on the phenotype can be variable on different cell lines or conditions.

To solve gene function variability problem, we introduced a new feature to quantify gene network property ^29^. This method utilized context-specific gene-gene interaction network and gene expression of neighborhood genes to calculate a gene network property score. We showed this score played an important role in the model prediction accuracy. Gene network property was also statistically negatively correlated with log2fc value, thus we believe that it can explain the impact of gene knockout on the phenotype. This finding has multiple implications on the application of CRISPR-Cas9 techniques. Technically, it is possible for us to enlarge the amount of available data to train machine learning and deep learning model. Most importantly, it suggests that the on-target efficiency and off-target effect determined in one condition may not be directly transfer to another condition unless the context-specific gene function is taken into account. It is critical when developing the CRISPR-Cas9 technique as a gene therapy.

Compared with DeepCpf1 and deepCrispr, seqCrispr performs better due to three reasons. 1) It involves a sequence feature engineering layer. It utilizes unsupervised representation learning to find the vector representation of 3mer instead of one-hot encoder. 2) We have investigated more biological features. The feature importance study implies that biological features can contribute 15%-20% to the model performance. 3) This hybrid model can take both advantages of RNN and CNN and make the model more resistant to data noise. However, there are still rooms to improve the performance. We believe that a better encoder of nucleotides can enhance model robustness and performance. Gene network property score can improve the way to display gene global function differences. More importantly, we used input permutation method to study features importance. It can be applied to any deep learning model and is shown to draw consistent conclusion. Thus, we could explore more advanced deep learning model. Even though next generation sequencing enables us to collect large-scale dataset, the amount of data is still less than sufficient. One issue is that the CRISPR-Cas9 experiment in each cell line lacks replication. Some cell lines have replication data but the experiments were conducted by different labs. Experiment performed on different lab will cause batch effect. We believed that more high quality data were still needed to reduce the noise of CRISPR-Cas9 experiment result and would improve the models performance.

## Methods

### Dataset

We have used three types of dataset. 1) Raw negative selection data. Data was collected from previously published literature ^28, 33^. We have picked up data from three cell lines K562, A549 and NB4. All three cell lines have the following features available: copy number variation, DNase-seq, Chip-seq for CTCF and H3K4me3 and RRBS data. Epigenetics features were collected as described in biological features section. Models were trained with normalized log2fc as model output. 2) Pre-selected negative selection data. These datasets were also used in other literature ^16^. Data were collected from three cell lines: HeLa, HL60 and HCT116. They have all biological features available except copy number variation. Data from negative selection dataset was preselected based on predicted gene essentiality. Only sgRNA targeting on predicted essential genes were selected. 3) CRISPR-Cpf1 dataset. The first two sets of data were from CRISPR-Cas9 system experiment, while this dataset was data from CRISPR-Cpf1 system experiment ^17^.

### Biological Features

Gene network property score integrated both global gene-gene interaction network information from STRING and gene expression data from Broad Institute Cancer Cell Line Encyclopedia (CCLE) ^13, 34^ or The Encyclopedia of DNA Element (ENCODE) ^13^. In this method, for each gene, its network property was calculated with the following steps: 1) Find genes which are connected with the studied gene 2) Calculate the product of connected gene expression values and gene-gene connection confidence score. Gene-gene connection confidence score is the weight of gene-gene interaction in gene-gene network 3) Sum all the products to get the score.

Copy number variation data was collected from CCLE ^34^. DNA-sgRNA binding melting temperatures are calculated using different thermodynamic tables from Sugimoto ‘96 ^35^. DNase peak, CTCF peak, RRBS peak and H3K4me3 peak data are curated from ENCODE ^13^. DNase peak, CTCF peak and H3K4me3 peak data chromosome location was annotated with hg38 reference genome. RRBS peak data chromosome location was referred with hg19 reference genome and was then overlifted to hg38 reference genome with CrossMap ^36^.

### Features importance study

We used random forest and boosted regress tree models implemented by h2o package, which treats categorical feature as one feature naturally. In this way, it is easier to interpret categorical features importance. In Random Forest model, we set number of trees to 200 and use default values for other parameters. In deep learning study, we used input perturbation methods to study the feature importance described here^37^. For each feature, we shuffled 40 times and use the average performance based on mean square error as feature importance. We then normalized each feature importance using the sum of all features importance. We implemented this method in python which is compatible with any deep learning model implemented in Keras.

### SeqCrispr model

The first part of SeqCrispr model is an embedding layer, in which 3mer’s representation was initialized by the vector learned with 3mer2vec and we made this vector representation trainable except in transfer learning. Each vector is the representation of a 3mer, who is the three nucleotides from position i to position i+2 on sgRNA 20 nucleotides. The last 3mer is a vector representation of PAM sequence. This was connected to a convolutional neural network, recurrent neural network or the combination of them. The last layer is a fully connected neural network. The input of fully connected neural network is the concatenation of all biological features and the output of the front layer. A linear regression was used to generate the final output. Mean square error was used as the loss function but spearman correlation was used as report performance metric. SeqCrispr model was implemented with Keras, which is built on top of Tensorflow. The implementation of SeqCrispr is available at https://github.com/qiaoliuhub/seqCrispr

## Acknowledgement

This work was supported by Grant Number R01LM011986 from the National Library of Medicine (NLM), Grant Number R01GM122845 from the National Institute of General Medical Sciences (NIGMS), and Grand Number R01AD057555 of National Institute of Aging of the National Institute of Health (NIH) as well as CUNY High Performance Computing Center.

## Author contributions

Q. L. and D. H. conducted the designed researches, analyzed the results, and wrote the paper. L. X. conceived the idea for the project, provided advice for experiment design, and revised the paper.

## Conflict of interest

The authors declare that they have no conflicts of interest with the contents of this article.

## Data Availability

The data that support the findings of this study are available from the corresponding author upon reasonable request.

