## Supplemental material for "Identifying Context-specific Network Features for CRISPR-Cas9 Targeting Efficiency Using Accurate and Interpretable Deep Neural Network"

**Supplementary information**

**sgRNA sequence encoding process**

To make sgRNA sequences suitable as input for classical convolutional neuron network, we designed a process that converts sgRNA sequences into 2-dimensional (or 3-dimensional) tensors. The process is described as follow,

We first split each sgRNA sequence of length $n$ into a list of $(n-2)$ 3-mers (3-nucleotide sub-sequences), for example, the sequence ACTGCG will be converted into {ACT, CTG, TGC, GCG}.

1. **3-mer numerical representations:**
   1.1. **One-hot encoding:** we used a 64-long one-hot vector to stand for each different 3-mers.
   1.2. **Word2vec embedding**: we used word2vec-learned embedding vectors as the representation for each 3-mer. Specifically, we converted all the human genome coding sequences into 3-mer format, and then use it as the corpus for the word2vec training.
2. **sgRNA sequence tensor representation:**
   2.1. **Vertical stacking**: We stacked all 3-mer numerical vectors for a sgRNA sequence vertically to construct 2-dimensional tensor. For example, sequence of length $n$ will be converted into a $\left( n-2 \right)\times64$ tensor if using one-hot 3-mer encoding.
   2.2. **Hilbert-curve filling**: inspired by [1], we adopted the similar idea as an alternative approach to align 3-mer numerical vectors into tensors. In this approach, we first used Hilbert curve filling to identify each 3-mer’s location in a 2-dimensional space, then extended the numerical representation of each 3-mers as the third dimension. As a result, each sgRNA sequence will be converted into a 3-dimensional tensor.

**SeqCrispr CNN-only performance:**

We compared SeqCrispr CNN-only models’ performance when useing different preprocessing over sgRNA sequences in three different cell lines. Specifically, the network model we tested consists of three convolutional layers (with max-pooling), and fully-connected layers in the end to obtain the prediction score.

**Table S1 Performance (Spearman correlation) comparison of SeqCrispr CNN-only model using different representations of nucleotides**

| **Preprocessing** | K562 | A549 | NB4 |
| --- | --- | --- | --- |
| One-hot encoding + vertical stacking | 0.42 | 0.38 | 0.37 |
| One-hot encoding + Hilbert-curve filling | 0.40 | 0.39 | 0.37 |
| Word2vec embedding + vertical stacking | **0.45*** | 0.39 | 0.38 |
| Word2vec embedding + Hilbert-curve filling | 0.43 | **0.43*** | **0.42*** |
